# Lassa virus NP DEDDh 3′-5′ exoribonuclease activity is required for optimal viral RNA replication and mutation control

**DOI:** 10.1101/2023.04.12.536665

**Authors:** Cheng Huang, Emily Mantlo, Slobodan Paessler

## Abstract

Lassa virus (LASV), a mammarenavirus from *Arenaviridae*, is the causative agent of Lassa fever (LF) endemic in West Africa. Currently, there are no vaccines or antivirals approved for LF. The RNA-dependent RNA polymerases (RdRp) of RNA viruses are error-prone. As a negative-sense RNA virus, how LASV copes with errors in RNA synthesis and ensures optimal RNA replication are not well elucidated. LASV nucleoprotein (NP) contains a DEDDH 3′-to-5′ exoribonuclease motif (ExoN), which is known to be essential for LASV evasion of the interferon response via its ability to degrade virus-derived double-stranded RNA. Herein, we present evidence that LASV NP ExoN has an additional function important for viral RNA replication. We rescued an ExoN-deficient LASV mutant (ExoN- rLASV) by using a reverse genetics system. Our data indicated that abrogation of NP ExoN led to impaired LASV growth and RNA replication in interferon-deficient cells as compared with wild-type rLASV. By utilizing PacBio Single Molecule, Real-Time (SMRT) long-read sequencing technology, we found that rLASV lacking ExoN activity was prone to producing aberrant viral genomic RNA with structural variations. In addition, NP ExoN deficiency enhanced LASV sensitivity to mutagenic nucleoside analogues in virus titration assay. Next-generation deep sequencing analysis showed increased single nucleotide substitution in ExoN- LASV RNA following mutagenic 5-flurouracil treatment. In conclusion, our study revealed that LASV NP ExoN is required for efficient viral RNA replication and mutation control. Among negative-sense RNA viruses, LASV NP is the first example that a viral protein, other than the RdRp, contributes to reduce errors in RNA replication and maintain genomic RNA integrity. These new findings promote our understanding of the basics of LASV infection and inform antiviral and vaccine development.

**Authors Summary:** Lassa fever (LF) is a severe and often fatal disease endemic in West Africa. There is no vaccines or antivirals approved for LF. The disease is caused by Lassa virus (LASV), a member of the arenavirus family. LASV nucleoprotein (NP) contains a DEDDh exoribonuclease (ExoN) motif, through which NP degrades virus-derived, immunostimulatory double-stranded RNA and inhibit host innate immune response. Thus, it is well known that NP ExoN is important for LASV pathogenicity. Intriguingly, the NP ExoN motif is highly conserved among arenaviruses, regardless of pathogenicity and viral ability to evade innate immune response, suggesting arenavirus NP ExoN may have additional function(s) in virus infection. In this study, we found that loss of ExoN activity affected LASV multiplication and RNA replication in interferon-deficient Vero cells. The ExoN-deficient rLASV exhibited reduced level of viral RNA, increased frequency of structural variation in virus genomic RNA, and higher mutation rate following mutagenic nucleoside analogue treatment. In conclusion, LASV NP ExoN plays an important role in viral RNA replication and fitness. Our new findings may inform antiviral and vaccine development and have broader implication on the function of NP ExoN of other arenaviruses.

## Introduction

The *Arenaviridae* family consists of four genera: *Mammarenavirus*, *Reptarenavirus*, *Hartmanivirus*, and *Antennavirus* (1, 2).*Mammarenaviruses*(referred to as arenaviruses hereafter) contain several pathogens of major clinical importance. Lassa virus (LASV) causes Lassa fever (LF), which is endemic in West Africa (3–6). The New World arenaviruses Junín virus (JUNV) and Machupo virus (MACV) cause Argentine hemorrhagic fever (AHF) and Bolivian hemorrhagic fever (BHF), respectively, in South America (7–10). These arenaviruses normally infect their rodent hosts, often persistently without overt disease signs. Spillover to humans occurs through aerosol inhalation and causes severe zoonotic diseases, of which vaccines and antivirals are limited (3, 8, 11–13). Accordingly, LASV, JUNV and MACV are classified as Category A Priority Pathogens in the USA. The World Health Organization has listed LF in the Blueprint list of priority diseases for which there is an urgent need for accelerated research and development (14).

Arenaviruses are negative-sense RNA viruses with a single-stranded, bi-segmented RNA genome (15). The large (L) segment RNA and the small (S) segment RNA are around 7.3 kb and 3.4 kb in length (Fig 1A). The L RNA encodes the RNA-dependent RNA polymerase (RdRp) L protein and a small, zinc finger protein (Z), which drives the assembly and budding of virus particles. The S RNA encodes the viral glycoprotein (GP), which mediates virus entry into host cells, as well as the nucleoprotein (NP), which is the major structural component of the nucleocapsid. On each genomic RNA, there is a highly structured, GC-rich intergenic region (IGR) (Fig 1A), which is the termination signal for viral mRNA transcription and is also required for packaging of viral genomic RNA into progeny virus particles (16). After virus entry, NP mRNA and L mRNA are transcribed from the 3′-end of S genomic RNA (S gRNA) or L genomic RNA (L gRNA), respectively (Fig 1B). Later in infection, S anti-genomic RNA (S agRNA) and L anti-genomic RNA (L agRNA) are synthesized from S gRNA and L gRNA, which are the templates for GPC mRNA and Z mRNA transcription as well as S gRNA and L gRNA synthesis (Fig 1B).

**Figure 1.**
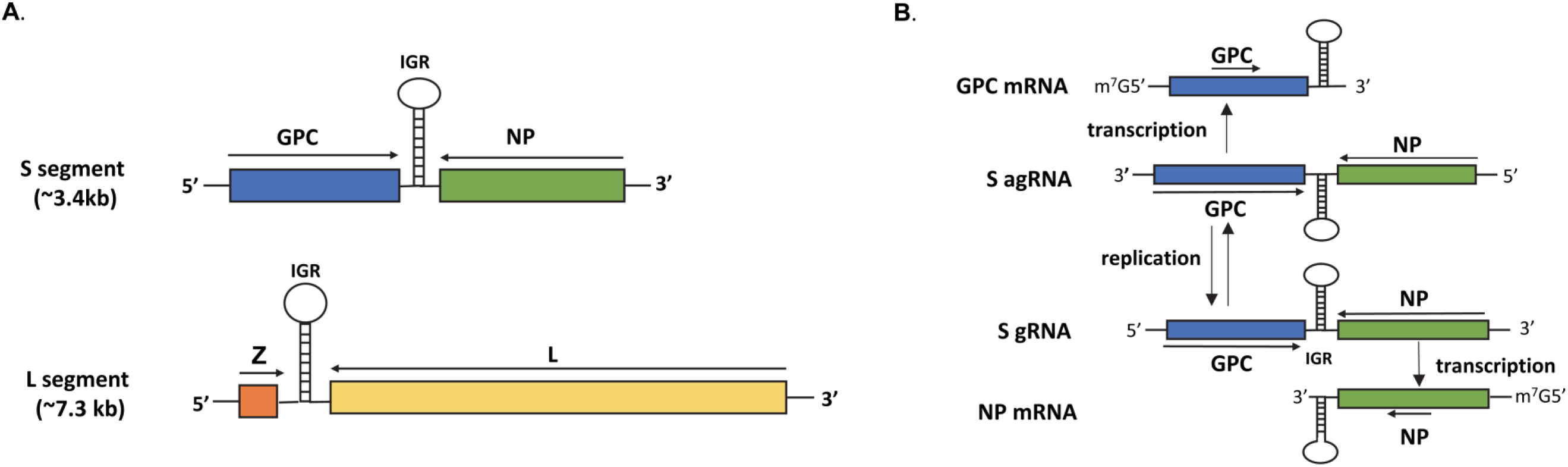
**(A).** Schematic diagram of arenavirus S and L genomic RNA. On each genomic RNA, an Intergenic Region (IGR) separates two ORFs. The highly structured IGRs are the termination signals of viral mRNA transcription and are required for efficient packaging of viral genomic RNA into progeny virions. **(B).** Genomic RNA replication and mRNA transcription of arenavirus S RNA. After virus entry, L protein and NP transcribe NP mRNA from S gRNA. From S gRNA, S antigenomic RNA (S agRNA) is also synthesized, which is the template for GPC mRNA and S gRNA synthesis later in virus infection. Arenavirus mRNA lacks a poly-A tail and contains the structured IGR at the 3′-end.

The RNA polymerase of RNA virus lacks proofreading activity. As negative sense RNA viruses, how arenaviruses control errors in RNA replication are not well elucidated. Both RdRp L protein and NP are required for viral RNA replication (17, 18), which occurs in virus-induced, discrete cytosolic structures (19). Arenavirus NP contains two structurally and functionally separated domains. The N-terminal half (aa 1-340 in LASV NP) binds to viral genomic RNA to form vRNP (20). In the C-terminal half of LASV NP, D389, E391, D466, D533 and H528 constitute a DEDDh 3′-5′ exoribonuclease (ExoN)- like motif. Structural and biochemical studies have established that the LASV NP harbors exoribonuclease activity specific for double-stranded RNA (dsRNA). Mutation of any of these DEDDh residues abolishes the ExoN activity (21–23). NP ExoN-mediated dsRNA degradation is essential for LASV NP inhibition of the Sendai virus-induced interferon response (21, 23), which is consistent with LASV suppression of the innate immune response *in vitro* and *in vivo.* It is widely accepted that LASV NP ExoN is critical for LASV immune evasion and pathogenicity (24, 25).

Interestingly, the NP DEDDh motif and its 3′-5′ ExoN activity are highly conserved in arenaviruses regardless of pathogenicity and virus ability to evade IFN response. For instance, the ExoN motif is also found in the NP of the non-pathogenic Tacaribe virus (TCRV), Pichinde virus (PICV) and Mopeia virus (MOPV) (22, 26–30). Notably, it has been shown that abrogation of ExoN activity impaired the ability of NP to support LASV minigenome replication (31, 32), suggesting a role of NP ExoN in arenavirus lifecycle aside from immune evasion. Interestingly, the nsp14 protein of the positive-sense RNA coronavirus also possesses the same DEDDh ExoN motif, which is essential for the fidelity of coronavirus RNA replication (33–35). Thus, further studies are warranted to investigate whether LASV NP ExoN plays a role in viral RNA replication in addition to immune evasion.

In the present study, we rescued an ExoN-deficient mutant LASV (ExoN- rLASV) and found that loss of ExoN activity resulted in impaired virus growth and viral RNA replication in interferon-deficient cells. Our study also showed that abrogation of NP ExoN led to increased production of aberrant LASV RNA and higher sensitivity to mutagenic nucleoside analogues as compared with wild-type LASV. These data present evidence that LASV NP ExoN has an important function in viral RNA replication in addition to its role in immune evasion. Our new findings may enhance our understanding of the basic virology of this important human pathogen and open new directions for future studies.

## Results

### Abrogation of NP ExoN activity impaired LASV multiplication in interferon-deficient cells

In biochemical studies, mutation of any of the LASV NP D389, E391, D466, D533 and H528 residues to alanine diminished the ExoN activity. Accordingly, we attempted to rescue recombinant LASV (rLASV, Josiah strain) harboring NP D389A and NP D389AG392A mutations to investigate the role of LASV ExoN in infection. To minimize the likelihood of reversion, two nucleotide substitutions were introduced to mutate the NP D380 and NP G392 residues. Specifically, for the NP D380A mutant, nt GA_1266_C_1267_ in S RNA (in antigenomic sense) was mutated to GC_1266_T_1267_. For the NP D389AG392A mutant, GA_1266_C_1267_ (D389) and GG_1275_A_1276_ (G392) in S RNA were mutated to GC_1266_T_1267_ (D389A) and GC_1275_T_1276_ (G392A), respectively. We successfully rescued the rLASV NP D389A mutant (ExoN- rLASV hereafter) by using the reverse genetics systems established in our lab. Sequencing analysis confirmed that the mutation is stable after 3 passages in Vero cells (Fig 2A). We could not rescue the rLASV NP D389AG392A mutant despite repeated attempts, indicating rLASV with the NP D389AG392A mutation is not viable.

**Figure 2.**
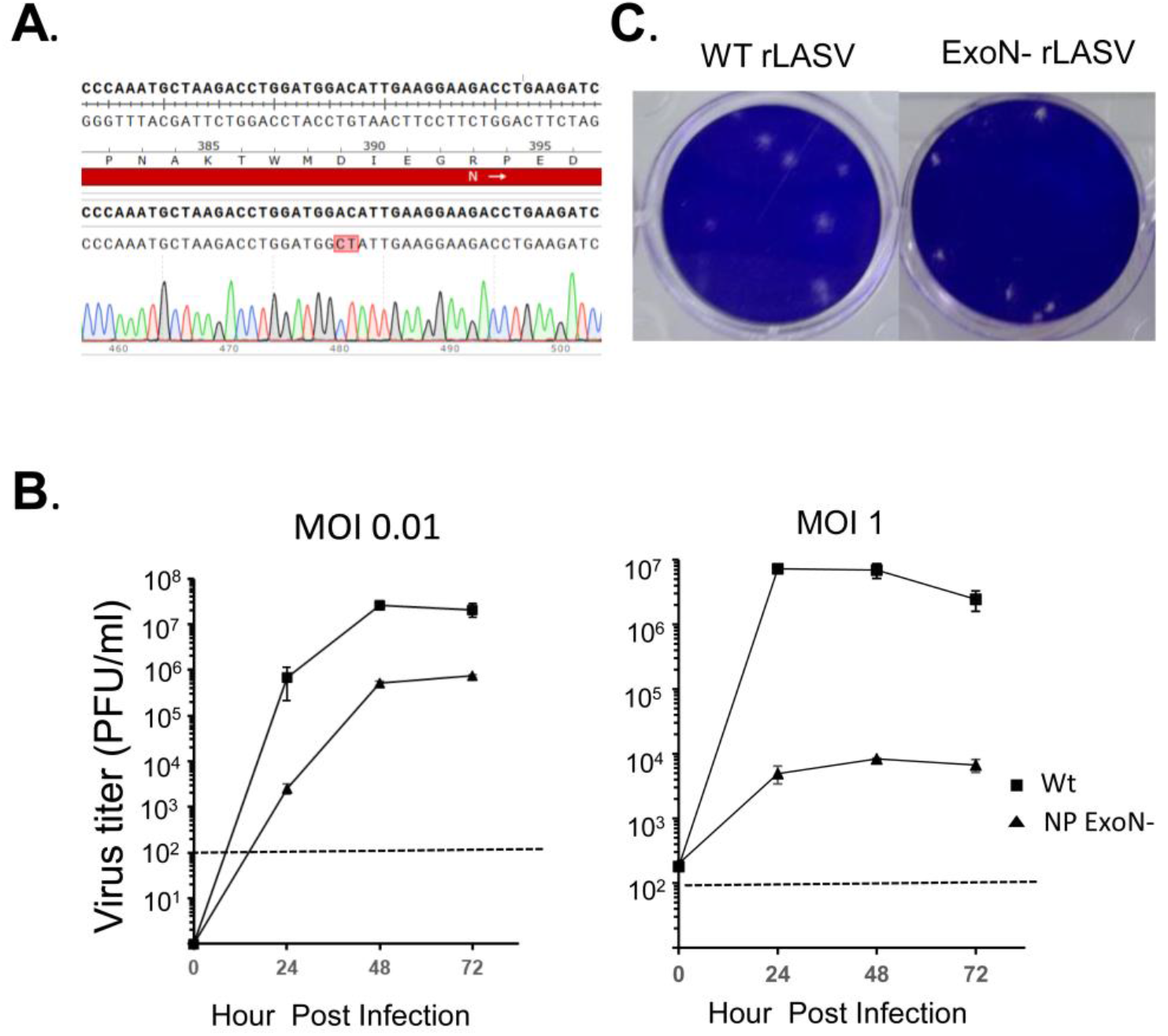
Abrogation of NP ExoN activity impaired LASV multiplication in interferon-deficient cells. A recombinant LASV (rLASV) with NP D389A mutation was rescued (ExoN- rLASV). (**A**). The D389A mutation was maintained after 3 passages in Vero cells (Sanger sequencing). (**B**). The growth curve of wild-type rLASV (wt) and the ExoN- rLASV (NP ExoN-) in Vero cells at multiplicity of infection (MOI) 0.01 and MOI 1. Dashed lines denote the detection limit of plaque assay. (**C**) The plaque morphology of wt rLASV (wt) and ExoN- rLASV.

We assessed the one-step and multiple-step growth kinetics of ExoN- rLASV in interferon-deficient Vero cells. The multiplication of ExoN- rLASV was attenuated at both conditions as compared with wild-type (wt) rLASV (Fig 2B). At 48 hours post-infection (hpi), the titer of ExoN- rLASV was 835-fold lower than wt rLASV at MOI 1, and 51-fold lower at MOI 0.01. Consistently, ExoN- rLASV formed smaller plaques in Vero cells as compared with wt rLASV (Fig 2C). These results indicated that abrogation of ExoN activity impaired rLASV propagation in Vero cells.

### NP ExoN deficiency impaired LASV RNA replication in interferon-deficient cells

As Vero cells are IFN-deficient (36, 37), the attenuated growth of ExoN- rLASV in Vero cells suggests that NP ExoN plays an important role in LASV replication other than viral evasion of the IFN response. We reasoned that ExoN deficiency may affect LASV RNA replication like its coronavirus nsp14 counterpart and further examined LASV RNA replication in Vero cells at conditions representing one-step virus growth (MOI 1 for 24 hr) and multi-step growth (MOI 0.01 for 72 hr). As shown in Fig 3A, the titer of ExoN-rLASV was 6671-fold lower than that of wt rLASV at MOI 1 and was 51.4-fold lower at MOI 0.01. We purified total RNA from infected cells and performed reverse transcription with random primers. Then, we conducted a qPCR assay to quantify the viral RNA level at the NP locus, which measures the total RNA level of S gRNA, S agRNA and NP mRNA (Fig 3B). Similarly, we also determined the RNA level of L gRNA, L agRNA and L mRNA with a qPCR assay using primers specific for the L locus (Fig 3C). At a MOI of 1, which represents one-step virus growth, the RNA level of ExoN- rLASV RNA was 1637-fold lower than that of wt rLASV at the NP locus (P<0.01, student t test), and 122- fold lower (P<0.01) at the L locus. At a MOI of 0.01, which represents multi-step virus growth, the RNA level of ExoN- rLASV was 45.5-fold lower than that of wt rLASV at the NP locus (P<0.01, student t test), and 25.2-fold lower at the L locus (P=0.04, student t test).

**Figure 3.**
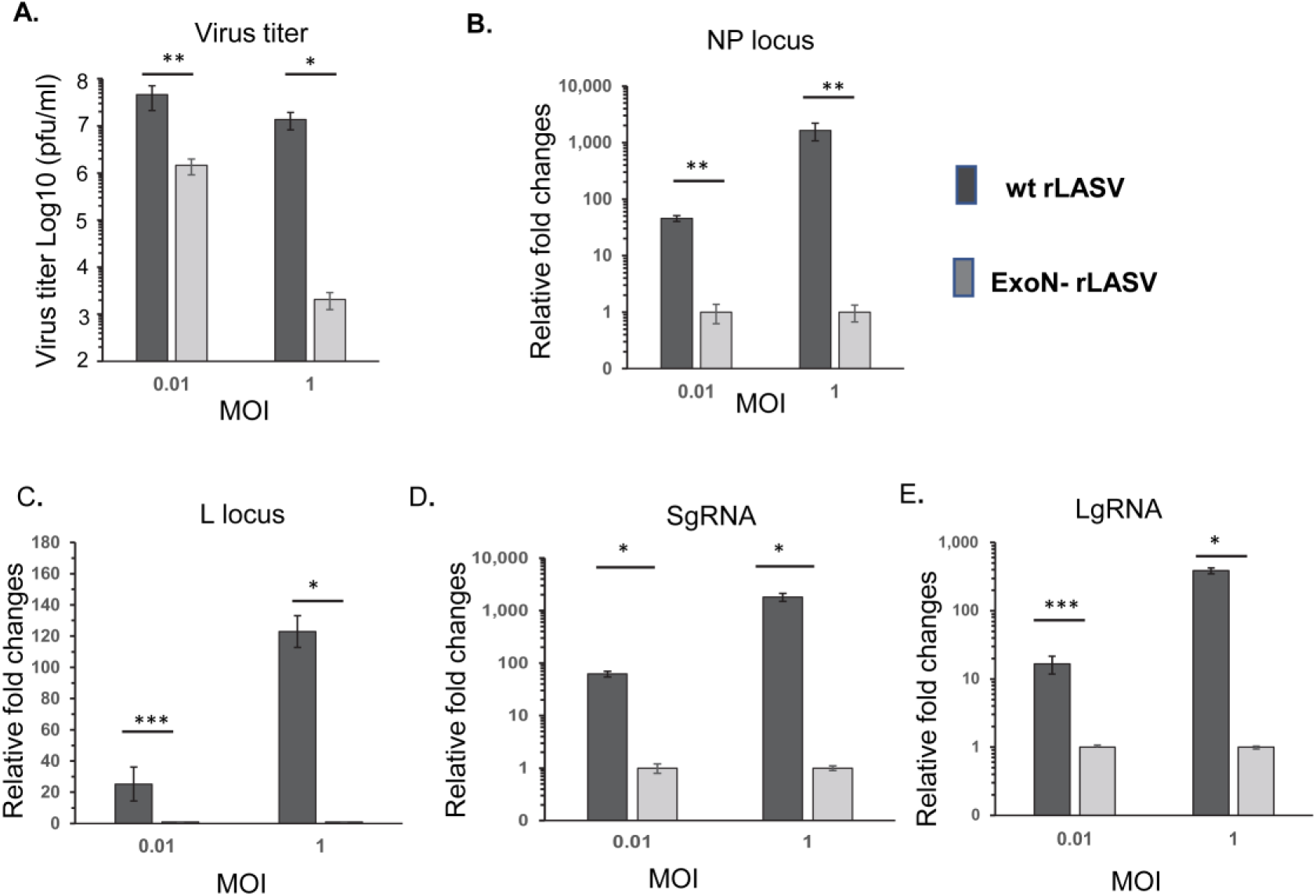
Abrogation of NP ExoN activity affected LASV RNA replication. Vero cells were infected with wt rLASV and ExoN- rLASV (MOI=0.01 and 1.0). At 72 hpi (MOI 0.01) or 24 hpi (MOI 1.0), virus titer was determined by plaque assay **(A)**. RT-qPCR assay for viral RNA level at the NP locus **(B)** and L locus **(C)**. The viral RNA level for S gRNA **(D)** and L gRNA **(E)** were also determined by qRT-PCR. The viral RNA level was normalized to the level of host β-actin mRNA and are presented as the fold changes relative to the level of ExoN- rLASV samples (set as 1.0). The data presents the mean and SEM of three independent experiments. (*: P<0.01; **: P<0.02; ***: P<0.05 with Student *t*-test).

We also examined the impact of ExoN deficiency on the level of S and L genomic RNA by RT-qPCR assay. In this assay, reverse transcription was performed with primers targeting the 3′-end of S genomic RNA and L genomic RNA using the high-fidelity reverse transcriptase SuperScript IV (Invitrogen), followed by qPCR assays targeting the NP locus on S gRNA and the L locus on L gRNA. At the one-step growth condition (MOI of 1), the level of ExoN- rLASV S gRNA was 1181-fold lower (P<0.01, t test) than that of wt rLASV; meanwhile the level of L gRNA was 387-fold lower (P<0.02) (Fig 3D and 3E). At a MOI of 0.01, the level of ExoN- rLASV S gRNA was 62.3-fold lower (P<0.01, t test) than that of wt rLASV; meanwhile the level of L gRNA was16.7- fold lower (P<0.02) than that of wt rLASV. These results clearly demonstrated that abrogation of ExoN affected LASV RNA level in IFN-deficient Vero cells, to an extent largely comparable with virus titer reduction.

### Abrogation of ExoN led to increased aberrant viral RNA formation in rLASV infection

Next, we performed RT-PCR to examine S gRNA and L gRNA in ExoN- rLASV infection. Vero cells were infected with wt rLASV and ExoN- rLASV at a MOI of 0.01 for 72 hours or at a MOI of 1 for 24 hours. Intracellular RNA samples were purified and reverse transcribed to cDNA using primers targeting the 3′-ends of S gRNA or L gRNA. Nearly full-length S gRNA (3391 nt) and L gRNA (7265 nt) was amplified with the high-fidelity Platinum SuperFi DNA Polymerase (Invitrogen). In agarose electrophoresis, we observed an aberrant S RNA product (indicated as S’ in Fig 4A) migrating slightly faster than the standard S gRNA product (3391 nt) in ExoN- rLASV samples at both a MOI of 0.01 and 1. The aberrant S’ product was detected specifically in ExoN- rLASV infection in repeated experiments. We also observed several aberrant L gRNA products (3.5 kb to 6 kb) in ExoN- rLASV samples at both a MOI of 0.01 and 1 (Fig 4B). These results demonstrated increased formation of aberrant S gRNA and L gRNA in ExoN- rLASV infection.

**Figure 4.**
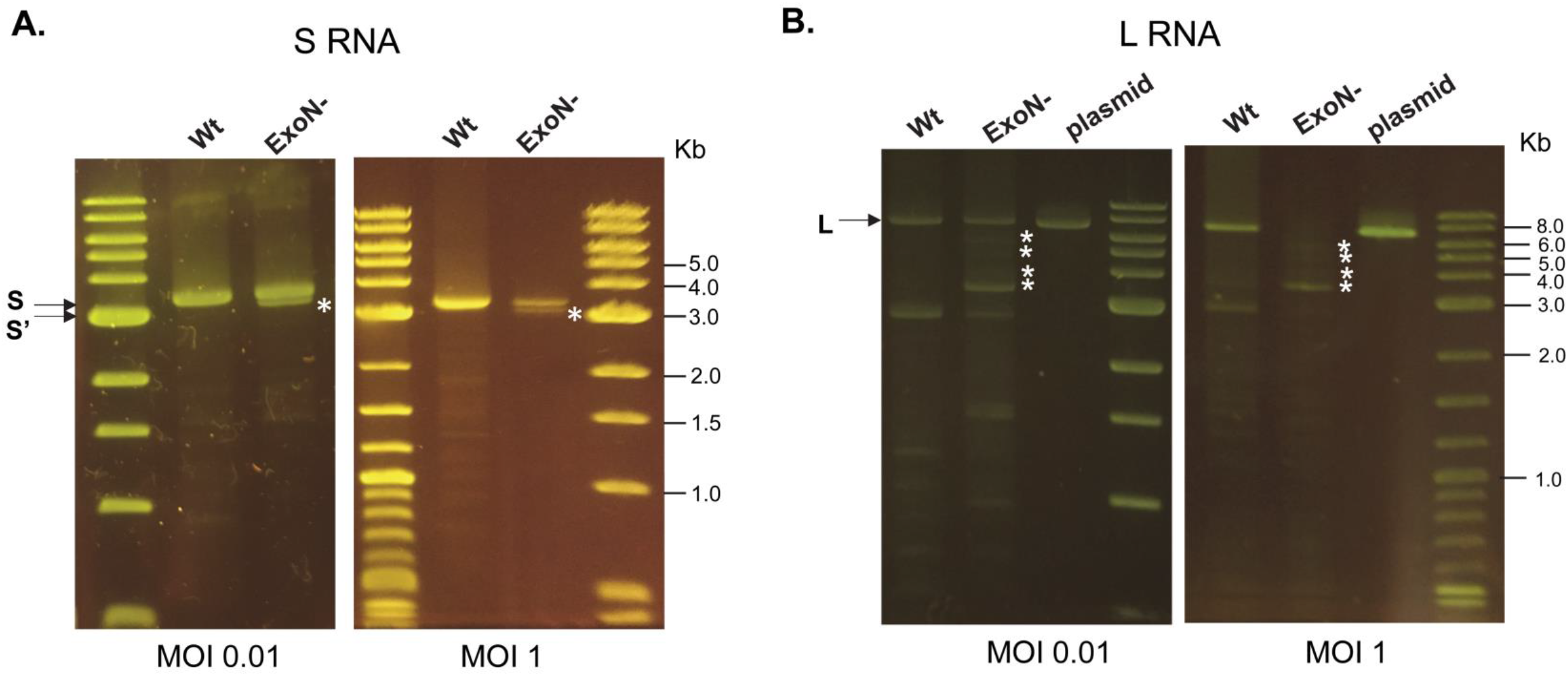
Aberrant viral RNA formation in ExoN- rLASV infection. Vero cells were infected with wt rLASV (wt) and ExoN- rLASV (ExoN-) at MOI 0.01 for 72 hr and at MOI 1 for 24 hr. RNA samples were purified from infected cells and transcribed to cDNA with primers targeting the 3′-end of S gRNA and L gRNA, respectively. High-fidelity PCR was performed to amplify close-to-full-length S gRNA (3391 nt) and L gRNA (7265 nt) followed by agarose gel electrophoresis. (**A**). RT-PCR result of S gRNA (S). S’ is a slightly fast-migrating aberrant S RNA product in ExoN- rLASV samples. (**B**). RT- PCR result of L gRNA. Several aberrant L gRNA products in ExoN- rLASV samples (3.5 kb to 6 kb) are observed. The aberrant RNA products are indicated by asterisk (*). Plasmid: PCR product using a plasmid containing LASV L segment as template.

### Abrogation of LASV NP ExoN activity affected the integrity of S gRNA

To determine the sequence of the aberrant S’ RNA in ExoN- rLASV infection, we performed agarose gel electrophoresis with ExoN- rLASV and wt rLASV S RNA PCR samples (MOI of 1) and purified the amplicons around the full size of S segment. Purified amplicons were cloned into a cloning vector (pSMART, Lucigen). Sanger sequencing of ten colonies identified two types of aberrant S RNAs formed in ExoN-rLASV infection. Three out of ten colonies had a 37-nt deletion (corresponding to nt1824-1860 in S agRNA), shown as the region from site I to site I’ in Fig 5A and 5B. The 37-nt deletion in the 67-nt-long IGR could disrupt the stem-loop structure of the IGR. In addition, five colonies contained a 231-nt deletion spanning the IGR and NP coding region (nt1616-1846 in S agRNA, from site II to II’ in Fig 5C). The 231-nt deletion consists of 42 nt in the IGR and 189 nt at the 3′ end of the NP gene (Fig 5C), which could disrupt IGR structure and cause an open reading frame shift from NP residue 507. In comparison, only two out of 10 colonies in the wt rLASV S samples had the 37-nt deletion in IGR, while the remaining colonies had the correct sequence. These data indicated that LASV lacking NP ExoN is more prone to forming structural deletion in S RNA, particularly around the IGR.

**Figure 5.**
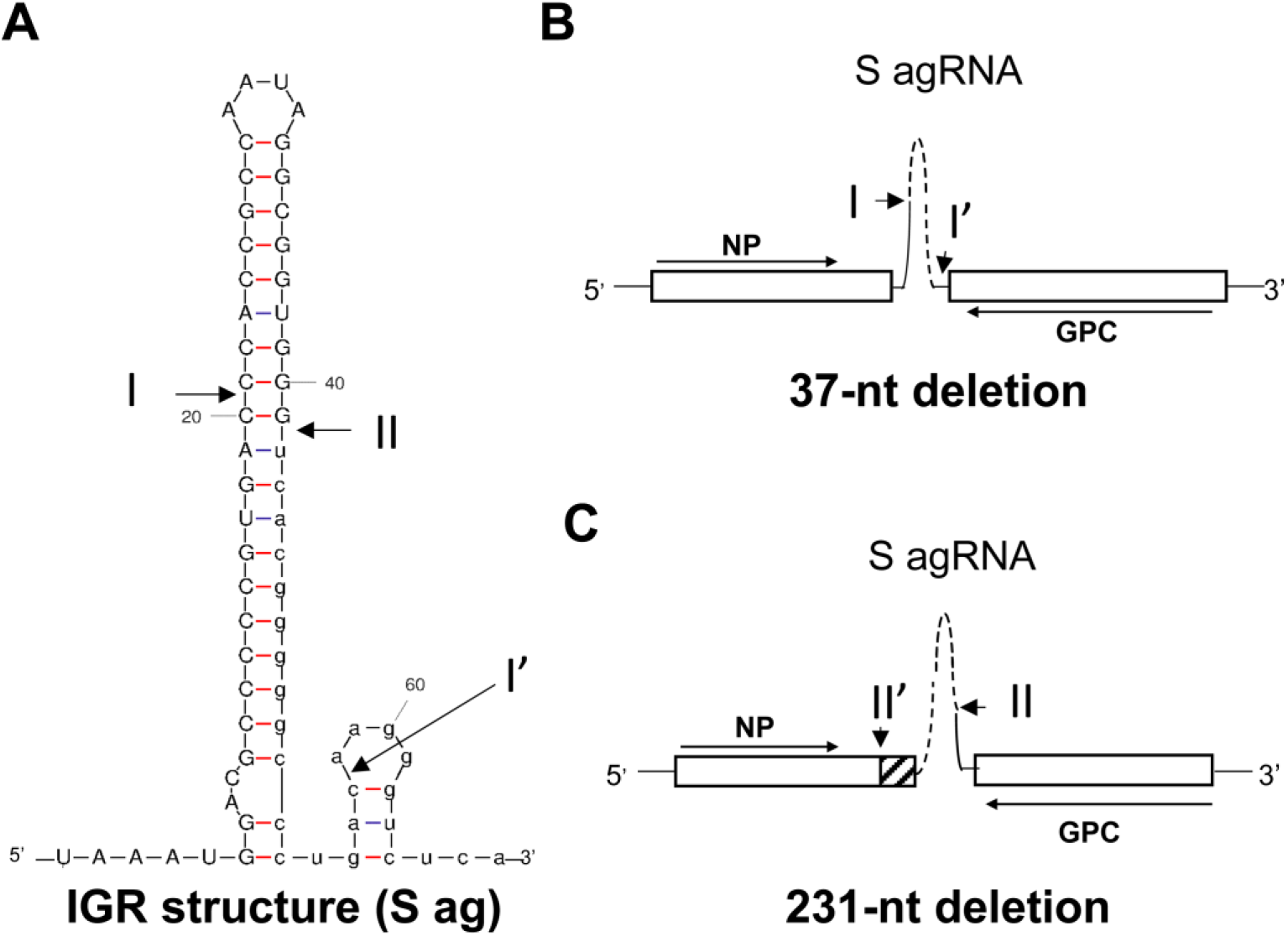
Sanger sequencing of aberrant S RNA formed in ExoN- rLASV infection. **(A)**. Predicted RNA structure of IGR of S RNA by *mFold.* **(B).** In ExoN- rLASV, a 37-nt deletion (nt1824-1860 in S agRNA) from site I to site I’ disrupted IGR structure. **(C)** a 231-nt deletion (nt1616-1846 in S agRNA) from the C-terminus of NP at site II’ to site II in the IGR (refer to A) was identified in the S gRNA sample of ExoN- rLASV.

### PacBio Single Molecule, Real-Time (SMRT) long-read sequencing to characterize aberrant S gRNA formed in ExoN- rLASV infection

To systematically assess the spectrum of aberrant S gRNA formed in ExoN- rLASV infection, we utilized PacBio Single Molecule, Real-Time (SMRT) long-read sequencing technology (38) to characterize LASV SgRNA at the single-molecule level. In PacBio SMRT long-read sequencing, a DNA polymerase continues to read a single circularized cDNA template for multiple rounds and provides deep sequencing data for each molecule. It can determine the circular consensus sequence (CCS) of full-length cDNA up to 10kb long at the single-molecule level with up to 99.99% accuracy. PacBio SMRT long-read sequencing is ideal for analyzing RNA quasi-species, long amplicons, and structural variations. It has been successfully used in studies on defective-interfering RNA in influenza virus infection and hepatitis C virus variants following drug treatment (39, 40).

We gel purified the nearly full-length S RNA PCR amplicon from wt- and ExoN- rLASV-samples (MOI of 1) and performed PacBio SMRT long-read sequencing (Sequel II). A total of 402,410 CCS reads and 381,134 CCS reads were obtained for wt rLASV and ExoN- rLASV samples, respectively (Fig 6A). The mean number of read passes for each cDNA was 28 times for wt rLASV (average length of 2319 nt) and 26 times for ExoN- rLASV (average length of 2610 nt) (Fig 6A). The mean read score of the assay was >0.999. CCS reads with a size of 3100-3391 bp were selected using Filter FASTQ and aligned with the LASV reference sequence using *minimap2* (41). Variants with higher than 3% frequency were called using *iVar* (42). In the ExoN- rLASV sample, we found that 88.9% and 4.9% of S RNA had the 231-nt and the 37-nt deletion, respectively (Fig 6B and 6C). In comparison, 7.1% and 18.2% of the S RNA in the wt rLASV sample had the 231-nt and the 37-nt deletion, respectively. The data also confirmed that greater than 99.93% of ExoN- rLASV harbored the intended NP D389A mutation. In addition, an A-to-G substitution at nt 393 (in antigenomic sense) was identified in S RNA of ExoN- rLASV with a frequency of 36.7%. A C-to-U substitution at nt 1518 in S RNA was found in wt rLASV sample with a frequency of 3.07%. Overall, a systematic analysis with PacBio SMRT sequencing clearly showed that LASV lacking ExoN had a higher frequency of structural variation in S gRNA.

**Figure 6.**
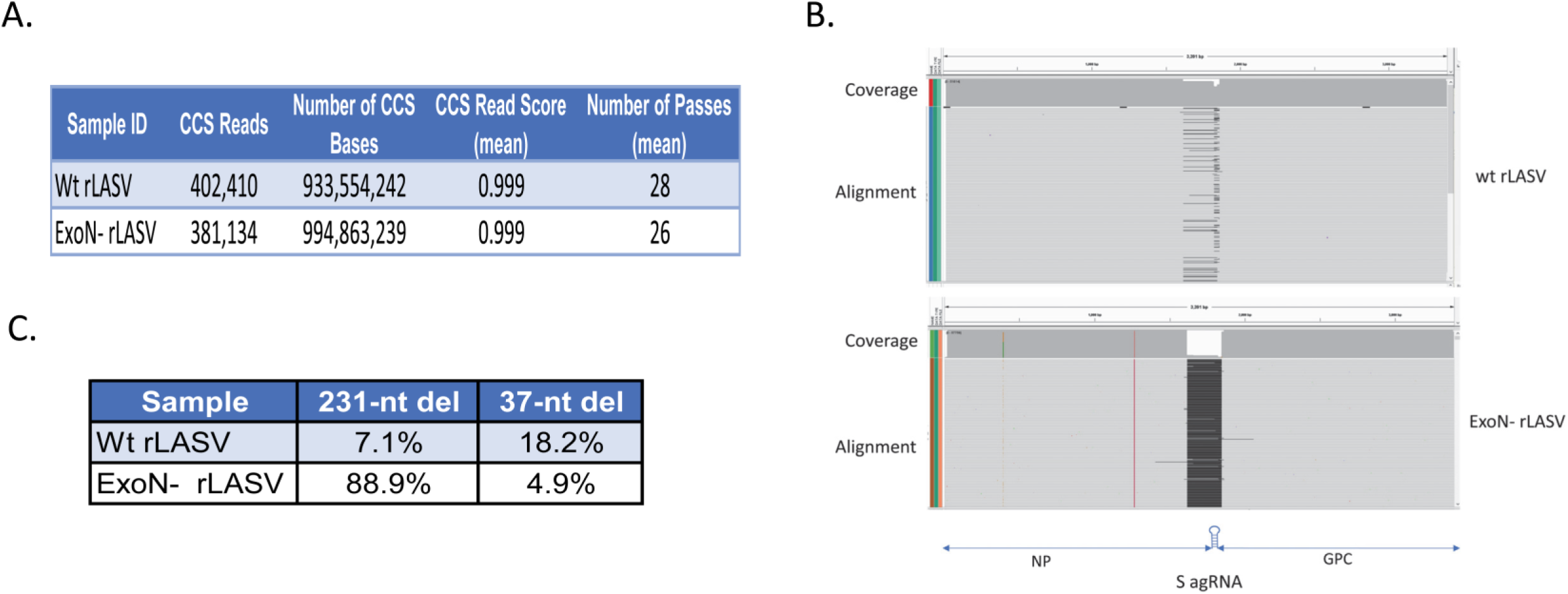
PacBio SMRT long-read sequencing of S gRNA in rLASV samples. Vero cells were infected with wt rLASV and ExoN- rLASV (MOI 1.0). RT-PCR product of the close-to-full-length SgRNA (3391 nt) were purified after agarose gel electrophoresis and subjected to PacBio SMRT long-read sequencing (Sequel IIe) with v3.0 chemistry. **(A).** HiFi CCS reads were generated using PacBio SMRTLINK v.10.1. **(B).** CCS reads with size 3100-3391bp were selected with Filter FASTQ and mapped to LASV reference seq (GenBank MH358389) with *minimap2.* Variants were called with *ivar* and visualized using Integrated Genomics Viewer. **(C)**. The percentage of the 231-nt deletion and the 37-nt deletion in S RNA of wt rLASV and ExoN- rLASV sample.

### ExoN- rLASV was more sensitive to nucleoside analogue treatment

Coronavirus nsp14 DEDDh 3′-5′ ExoN proofreads newly synthesized viral RNA by removing mis-incorporated nucleotides. Abrogation of nsp14 ExoN activity renders coronaviruses susceptible to lethal mutagenesis when treated with nucleoside analogues such as 5-fluorouracil (5-FU) (43). 5-FU treatment also increases the mutation rate of the prototype arenavirus lymphocytic choriomeningitis virus (LCMV) and affects viral fitness (44). To investigate if loss of ExoN activity could increase LASV sensitivity to mutagenic nucleoside analogues, we evaluated the sensitivity of wt rLASV and ExoN- rLASV to 5-FU in Vero cells (MOI of 0.1). 5-FU treatment alone did not affect cell viability at the concentrations administrated in this experiment (Fig. 7A, CellTiter-Glo Viability Assay, Promega). Abrogation of NP ExoN increased LASV sensitivity to 5-FU treatment starting from 100 µM (Fig 7B). At 400 µM, the virus titer of ExoN- rLASV deceased by 1053-fold as compared with mock treatment, which was 13.5-times greater than the 78-fold decrease for wt rLASV (Fig 7B).

**Figure 7.**
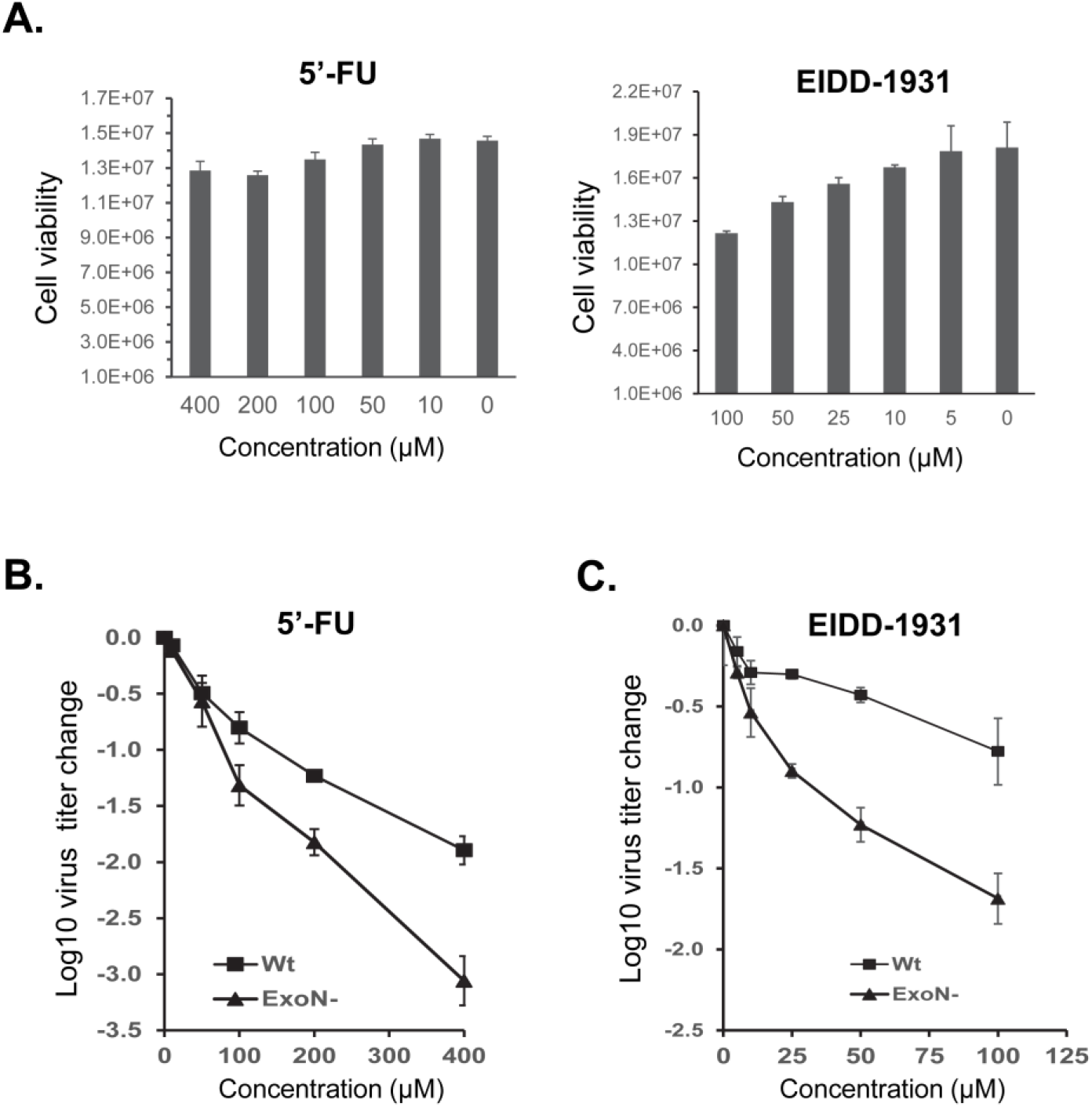
NP ExoN deficiency enhanced LASV sensitivity to mutagenic nucleotide analogues. **(A).** Vero cells were treated with 5′-FU and EIDD-1931 at indicated concentrations. At 48 hr post treatment, cell viability was measured using CellTiter-Glo Viability Assay (Promega). Data shown are the average (n=4) and SEM. **(B) and (C).** Vero cells were treated with 5-FU and EIDD-1931 at different concentrations as indicated. Cells were infected with wt rLASV (wt) and ExoN- rLASV (ExoN-) at MOI 0.1. At 48 hpi, virus titers were determined by plaque assay. Log10 virus titer changes relative to the virus titer of non-treated cells are shown. Data presented are the mean and the SEM of three independent experiments.

EIDD-1931 (*N4*-hydroxycytidine or NHC) is a cytidine analogue that increases the mutation frequency of a broad range of RNA viruses (45). We also assessed the sensitivity of ExoN- rLASV and wt rLASV to EIDD-1931. EIDD-1931 treatment alone (5 µM-100 µM) did not substantially affect cell viability (Fig 7A, CellTiter-Glo Viability Assay, Promega). The titer of ExoN- rLASV was decreased by 47.6-fold following 100 µM EIDD-1931 treatment, greater than the 6-fold reduction of wt rLASV at the same condition (Fig 7C). Collectively, these results indicated that abrogation of NP ExoN activity rendered LASV more sensitive to mutagenic nucleoside analogues.

### Increased rate of single nucleotide variation in ExoN- rLASV RNA following 5′-FU treatment

We further assessed the impact of the loss of ExoN on the mutation rate of LASV RNA following 5-FU treatment. We infected Vero cells with rLASV (MOI 0.1) and treated cells with a low level of 5-FU (100 μM). Extracted viral RNAs were reverse transcribed with primers specific to S genomic and L genomic RNA. We performed high-fidelity PCR to generate two overlapping amplicons (1.8 kb and 1.9 kb) for the S segment and three overlapping amplicons (2.5 kb, 2.6 kb and 3.2 kb) for the L segment. The amplicons were purified from agarose gels and subjected to Illumina next generation sequencing. The read depth was at least 1,000,000 reads at each site in the LASV genome (S1 Fig). Alternate alleles with variation frequency above 0.1% at each position of the genomic RNA were included for analysis.

We examined the distribution and the number of variants in viral genomic RNA from mock- and 5-FU-treated samples (Fig 8 and Table). In mock samples, a total of 87 sites in ExoN- LASV genomic RNA had alternate alleles with frequencies above 0.1%, higher than the 28 sites with alternate alleles in wt rLASV genomic RNA (Table). Among the 87 sites identified in ExoN- LASV genomic RNA, 65 sites were in L gRNA (Table). The low frequency of nucleotide substitution in 5-FU samples (0.1-1%) was correlated with the low level of 5-FU treatment (100 μM). At 100 μM 5-FU, 323 sites in ExoN- LASV genomic RNA had alternate alleles (Table), substantially higher than the 104 positions identified in wt LASV RNA.

**Figure 8.**
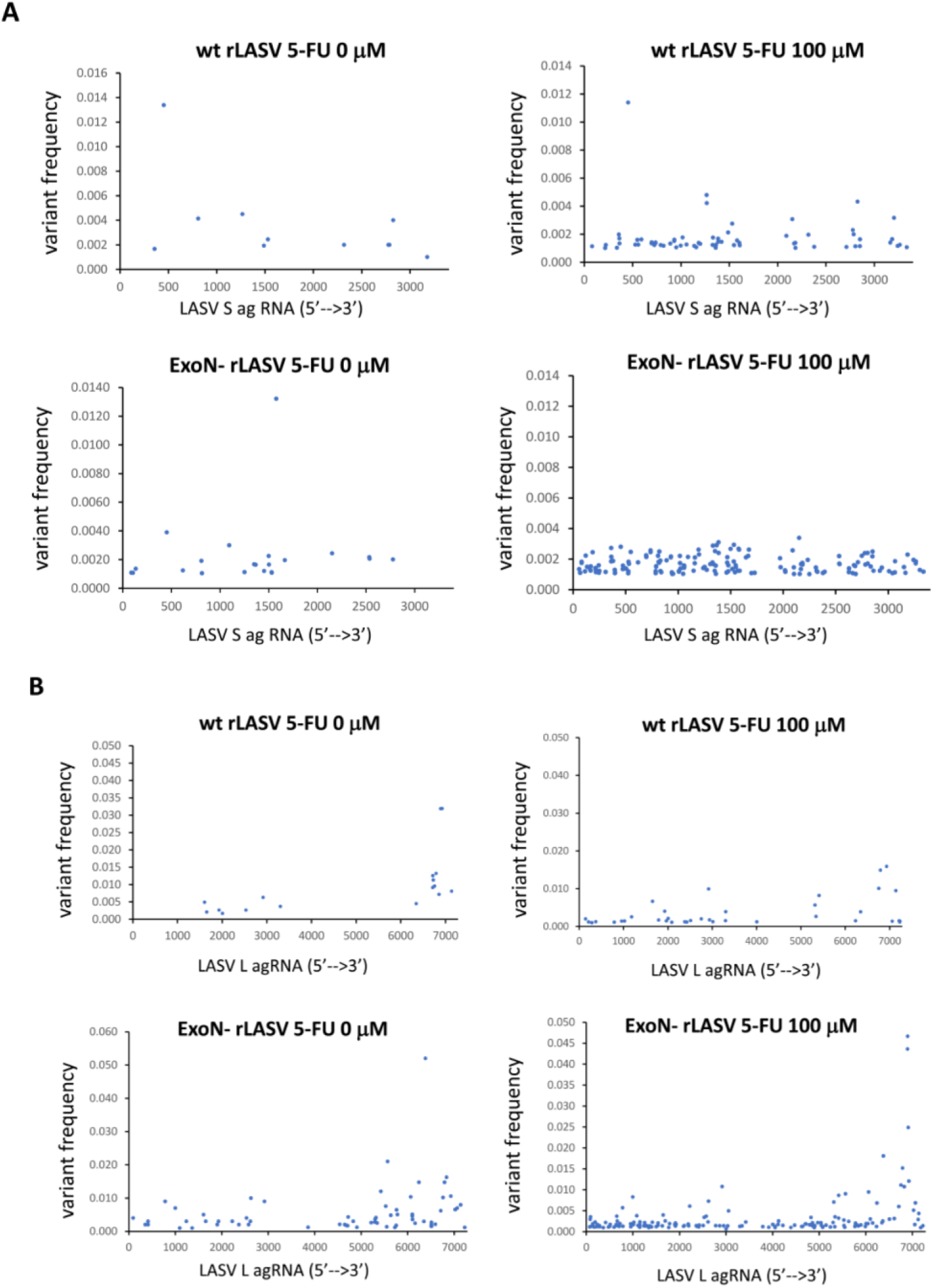
Distribution of alternate alleles in LASV genomic RNA following 5-FU treatment. Vero cells were mock-treated or treated with 5-FU and infected with wt rLASV and ExoN- rLASV (MOI 0.1). The distribution of alternate alleles in LASV S segment RNA **(A)** and L segment RNA **(B)** are shown. Y axis shows the frequency of variations on each position of viral genomic RNA. X axis shows the genomic position of LASV agRNA.

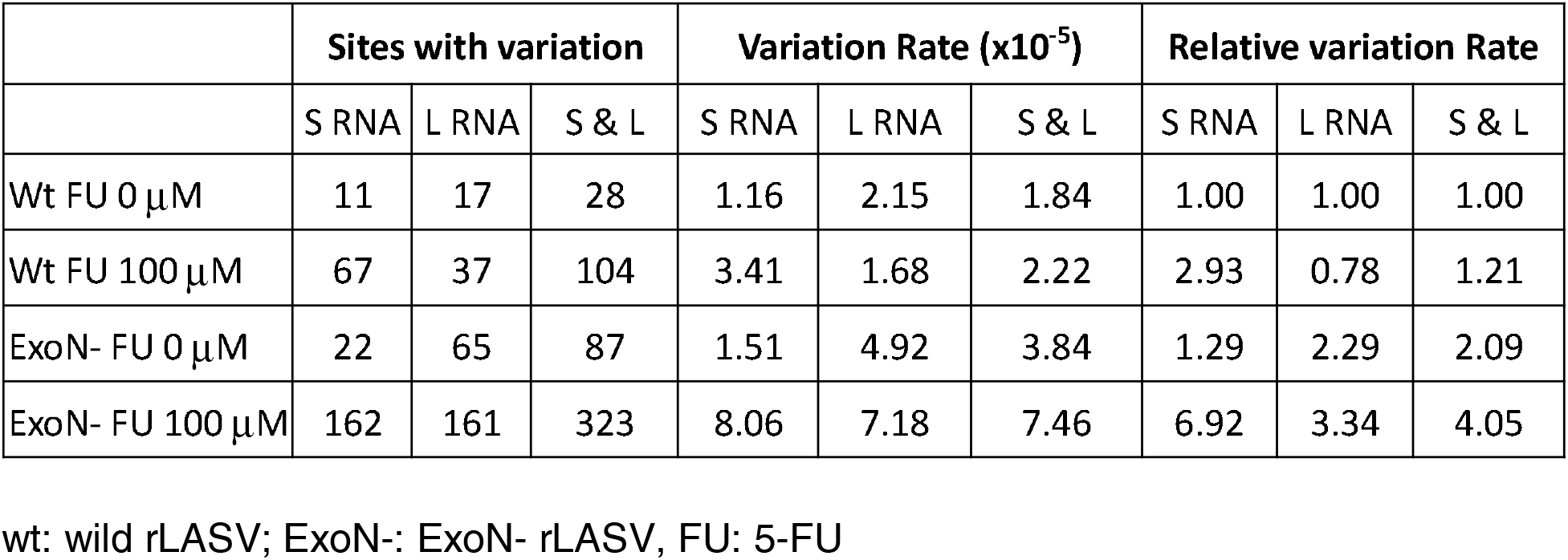
Table: NGS data of the number of sites and the variation rate of LASV genomic RNA following 5-FU treatment

| | Sites with variation | | | Variation Rate ($\times 10^{-5}$) | | | Relative variation Rate | | |
| --- | --- | --- | --- | --- | --- | --- | --- | --- | --- |
|  | S RNA | L RNA | S & L | S RNA | L RNA | S & L | S RNA | L RNA | S & L |
| Wt FU 0 μM | 11 | 17 | 28 | 1.16 | 2.15 | 1.84 | 1.00 | 1.00 | 1.00 |
| Wt FU 100 μM | 67 | 37 | 104 | 3.41 | 1.68 | 2.22 | 2.93 | 0.78 | 1.21 |
| ExoN- FU 0 μM | 22 | 65 | 87 | 1.51 | 4.92 | 3.84 | 1.29 | 2.29 | 2.09 |
| ExoN- FU 100 μM | 162 | 161 | 323 | 8.06 | 7.18 | 7.46 | 6.92 | 3.34 | 4.05 |

wt: wild rLASV; ExoN-: ExoN- rLASV, FU: 5-FU

We further compared the nucleotide variation rate of wt- and ExoN- rLASV genomic RNA (Table). In the absence of 5-FU, the variation frequency of wt rLASV genomic RNA was 1.84 x 10^-5^ per nucleotide read. Treatment with 100 μM of 5-FU increased the variation rate of wt rLASV genomic RNA by 21% to 2.22 x 10^-5^ per nt read. For ExoN- rLASV, the variation frequency of genomic RNA was 3.84 x 10^-5^ per nt read at 0 μM 5- FU, 2.09-fold of that of wt rLASV. At 100 μM 5-FU, the variation frequency of ExoN- rLASV genomic RNA was increased by 95% to 7.46 x 10^-5^ per nt read and was 3.34- fold of the variation frequency of wt rLASV.

5-FU incorporation causes A:G and U:C transitions in viral RNA during LCMV and SARS-CoV-1 infection (43, 44). In this study, we found A:G and U:C transitions at 78 sites (75.7%) and 278 sites (86.1%) in wt rLASV genomic RNA and ExoN- rLASV genomic RNA, respectively in 5-FU samples (Fig 9). In the mock samples, A:G and U:C transition was identified at 7 sites (25%) and 32 sites (36.8%) in wt rLASV and ExoN- rLASV genomic RNA, respectively.

**Fig 9:**
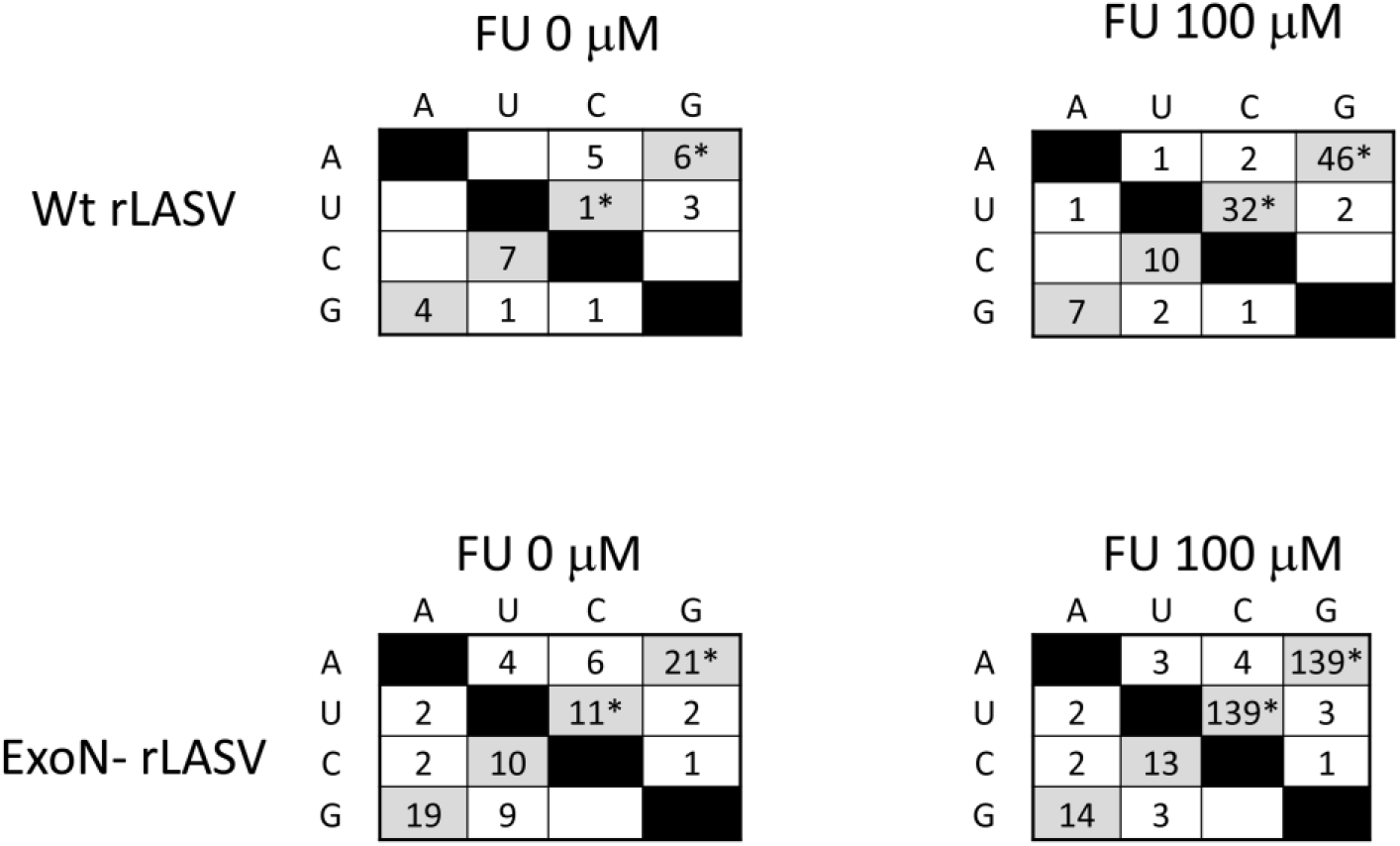
The number of positions with base changes in viral genomic RNA. All values represent the number of positions with alternate alleles at frequency higher than 0.1% in wt rLASV and ExoN- rLASV genomic RNA following 0 μM and 100 μM 5- FU treatment. Base transitions (A:G, G:A, U:C and C:U) are shaded in grey. 5-FU specific transitions (A:G and U:C) are marked with asterisk. Base transversions between purines and pyrimidines are shown in white.

In summary, the NGS data indicated that lack of ExoN activity increased the frequency of single nucleotide variation in LASV genomic RNA in both mock and sublethal 5′-FU conditions.

## Discussion

The LASV NP ExoN activity is known to be essential for LASV evasion of host innate immune responses. Interestingly, the DEDDh motif and the ExoN activity is highly conserved in the arenavirus family, regardless of viral pathogenicity, implying it has additional function(s) in arenavirus lifecycle. In this study, we found that abrogation of NP ExoN impaired LASV growth and RNA replication in IFN-deficient Vero cells. In addition, ExoN- rLASV was more prone to producing aberrant viral RNAs and more sensitive to mutagenic nucleoside analogues. Thus, LASV NP ExoN has a previously unrecognized function in LASV RNA replication, which reduces rates of viral RNA substitution and production of aberrant viral RNA. As the ExoN motif is highly conserved in arenaviruses, future studies should seek to determine whether the NP ExoN of other arenaviruses are also important in viral RNA replication.

In this study, we presented the evidence that abrogation of NP ExoN led to impaired LASV growth and smaller plaque size in IFN-deficient Vero cells (Fig 2). We also found that rLASV with NP D389AG392A mutation was not viable. These data demonstrated that NP ExoN is important for LASV fitness in addition to its role in immune evasion. Other groups have reported that deficiency in NP ExoN leads to attenuated growth for rMOPV, rPICV and the AV strain of rLASV (26, 27, 31). In addition, PICV possessing the NP D380A mutation (equivalent to the NP D389A mutation for LASV) forms smaller plaques in Vero cells. PICV wt revertant, which forms large plaques, could be readily identified after two passages of the NP D380A PICV mutant (27). Therefore, NP ExoN could be important for arenavirus fitness in general.

The present study provides evidence that abrogation of NP ExoN activity affected LASV RNA replication in IFN-deficient cells (Fig 3), which explains the impaired growth of ExoN- rLASV. For RNA viruses, the RNA replication is error-prone as RdRp lacks proofreading activity. When nucleotide mis-incorporation occurs, RdRp may pause or stop RNA elongation, which consequently affects viral RNA synthesis. Interestingly, MOPV NP and LCMV NP have been shown to excise mismatched nucleotides at the 3′- end of the dsRNA substrate in biochemical studies (28). Accordingly, it is plausible that LASV NP ExoN could also remove mis-incorporated nucleotides like MOPV NP and LCMV NP and facilitate viral RNA synthesis. In this scenario, lack of NP ExoN may affect the yield or the rate of arenaviral RNA synthesis, which may partly explain the impaired viral RNA replication in ExoN- rLASV infection. Further studies are required to investigate whether LASV NP ExoN can excise mismatched nucleotides, through which ExoN helps to reduce nucleotide mis-incorporation and facilitate viral RNA synthesis.

We detected increased production of aberrant S gRNA and L gRNA in ExoN- rLASV infected cells in repeated experiments (Fig 4). This data indicated that loss of LASV NP ExoN affected the integrity of viral genomic RNA. Sanger sequencing analysis of nearly full-length S gRNA amplicons identified two types of deletion mutants in ExoN- rLASV samples. Three of ten colonies had a 37-nt deletion (nt1824-1860, S agRNA) located in the 67-nt-long IGR (site I to site I′ shown in Fig 5A and 5B). Five colonies contained a 231-nt deletion (nt1616-1846 in S agRNA, from II to II’ shown in Fig 5C) in the IGR and NP coding region. The 37-nt deletion could disrupt the stem-loop structure in IGR, meanwhile the 231-nt deletion could disrupt the IGR structure and cause a shift in open reading frame from NP residue 507. In comparison, only two out of ten colonies of wt rLASV S samples had the 37-nt deletion in IGR. As the IGR is essential for transcription termination of arenavirus mRNA and packaging of genomic RNA into progeny virus particles, abrogation of NP ExoN may indirectly affect these two important steps in viral RNA replication.

To better understand the spectrum of LASV SgRNA variants, we utilized PacBio SMRT long-read sequencing to characterize LASV SgRNA amplicon at the single-molecule level. Short-read Illumina NGS has low sensitivity and high false positive rates in solving complex Structural Variations (SVs) with deletions and insertions at least 50 nt in size (46). PacBio SMRT long-read sequencing can determine the circular consensus sequence of full-length cDNA up to 10kb long at the single-molecule level, which is ideal for analyzing RNA quasi-species, long amplicons, and SVs. In addition, SMRT long-read technology is suitable for sequencing through highly repetitive, GC-rich sequences that are present in the IGRs of arenavirus RNA. In this study, raw reads with a minimum number of 3 passes and higher than 99% accuracy were used to generate the CCS reads to ensure read accuracy. The mean number of read passes for each cDNA was 28 times for wt rLASV with an average read length of 2319 nt, and 26 times for ExoN- rLASV with an average read length of 2610 nt (Fig 6A). The mean reading score of the SMRT assay was >99.9%. Variants with >3% frequency was selected in data analysis to ensure the reliability of the reads. The results demonstrated that 88.9% and 4.9% of the S genomic RNA contained the 231-nt deletion and the 37-nt deletion in ExoN- rLASV sample, respectively. In comparison, 7.1% and 18.2% of the S RNA in wt rLASV sample has the 231-nt deletion and the 37-nt deletion, respectively. In addition, higher than 99.93% of ExoN- rLASV S gRNA harbored the intended NP D389A mutation, which confirmed that the NP D389A mutation was maintained in ExoN- rLASV. Our data suggests that NP ExoN is important for viral control of structural deletion in LASV genomic RNA.

The molecular basis for these structural deletions formed in ExoN- rLASV S RNA is unclear. Sequence analysis revealed the presence of homologous sequences flanking the 37-nt deletion and the 231-nt deletion in S RNA (Fig 10, boxed sequences, sites I/I’ for the 37-nt deletion, sites II/II’ for the 231-nt deletion). Based on the homologous sequences, we propose a stop-and-realign model for the mechanism of increased structural deletions in S RNA in ExoN- rLASV infection (Fig 10). When RNA synthesis error occurs, LASV NP ExoN may excise the mismatched nucleotide, like its MOPV and LCMV counterparts (28), and allows the RdRp to resume RNA elongation. Lack of NP ExoN activity may lead to the stop of viral RNA synthesis, as shown in Fig 10A for the 37nt-deletion and in Fig 10B for the 231nt-deletion (Step 1, mis-incorporation and stop). Depending on the nucleotide mis-incorporated, LASV RdRp L protein may realign the 3′-end of nascent RNA (at site I in Fig 10A and site II in Fig 10B) with downstream homologous sequences on the template RNA (sites I’ and II’ in Fig 10A and Fig 10B, respectively) and resume RNA elongation (Step 2, realign and resume). As a result, RdRp skips the regions between these homologous sequences, leading to the 37-nt deletion (Fig 10A) and the 231-nt deletion (Fig 10B) in S RNA, respectively. Thus, LASV NP ExoN may control structural deletions by reducing errors in LASV RNA synthesis. In this regard, it is worth noting that RNA viruses with low-fidelity RdRp, such as Sindbis virus or tombusvirus, exhibit an increased production of defective viral genomic RNA correlated with an enhanced rate of viral RNA recombination (47–49). Further studies are warranted to investigate the molecular basis of increased aberrant RNA formation associated with LASV NP ExoN deficiency and the impact on virus infection.

**Figure 10.**
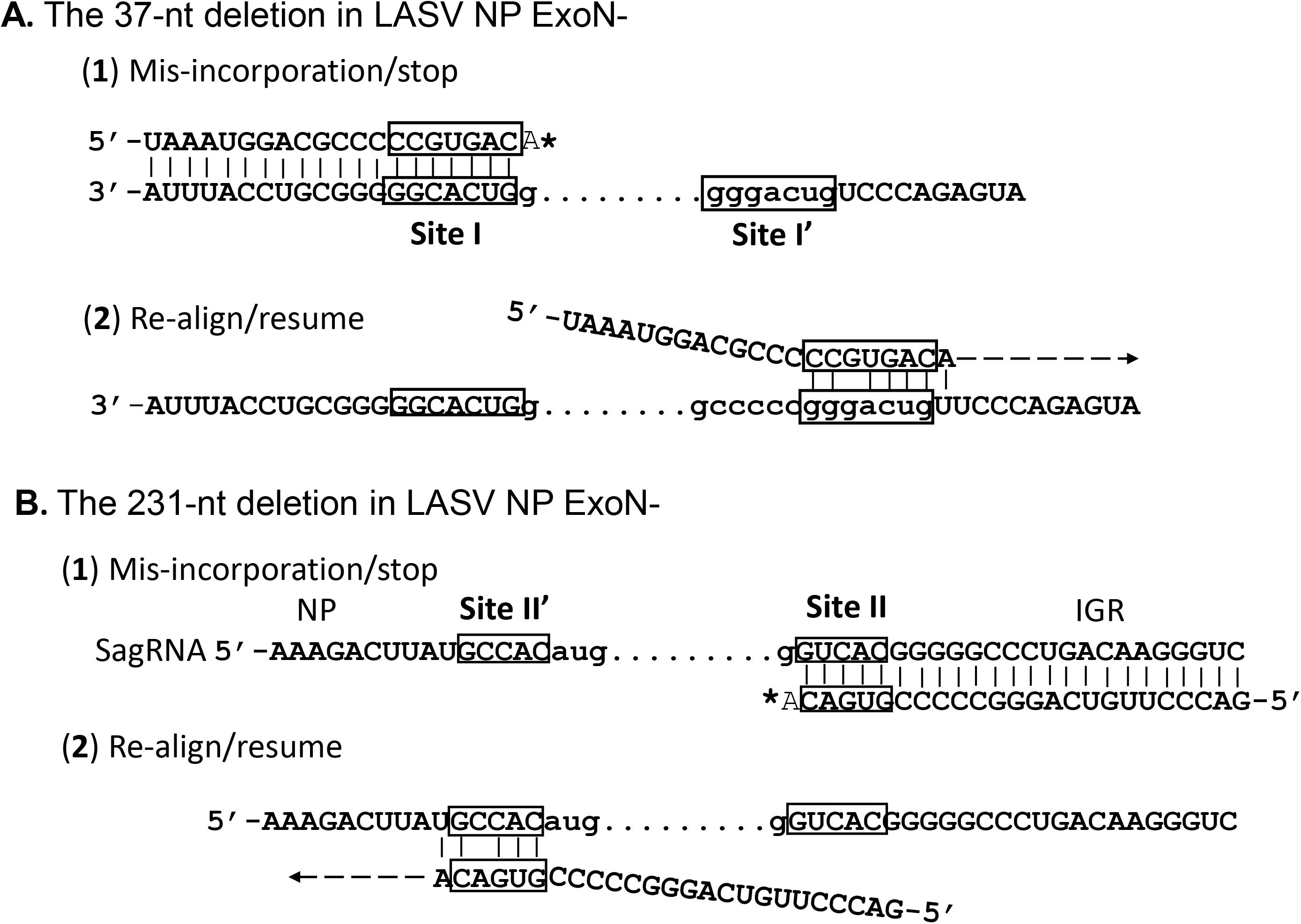
A stop-and-realign model for the 37-nt deletion (A) and the 231-nt deletion (B) in S RNA of ExoN- rLASV. Homologous sequences flanking the deletion sites are indicated as boxed sequences (sites I/I’ for the 37-nt deletion, sites II/II’ for the 231-nt deletion). Sites I and II are located in IGR stem-loop structure. In viral RNA synthesis, NP ExoN may remove mis- incorporated nucleotide and helps LASV RdRp L protein to resume viral RNA elongation. In the absence of NP ExoN, viral RNA elongation may terminate (Step 1). Depending on the nucleotide mis-incorporated (i.e., an A instead of C as shown in A and B), viral RdRp L protein may realign the 3′-end of nascent RNA (5′-CCGUGACA in site I or 5′-GUGACA in site II) with downstream homologous sequences on the template RNA (3′-GGGACUGU-5′ in site I’ and 3′-CACCGU-5′ in site II’) and resume RNA elongation (Step 2). Thus, the 37-nt deletion and the 231-nt deletion in S RNA may result from viral RdRp skipping the regions between the homologous sequences, an error that could be controlled by functional NP ExoN.

Our data of nucleoside analogue study support that LASV NP ExoN contributes to control the frequency of nucleotide substitution in viral RNA. In LCMV infection, 5-FU could be incorporated into nascent viral RNA and cause mutations that affect virus fitness. Consistently, abrogation of NP ExoN rendered LASV more sensitive to the mutagenic nucleoside analogues 5-FU and EIDD-1931 in virus titration assay (Fig 7). Our NGS data showed that ExoN deficiency is associated with increased nucleotide substitution in rLASV RNA following 5-FU treatment. At 100 μM 5-FU, the mutation rate of ExoN- rLASV was increased by 94%, while that of wt rLASV was increased by 21%. In the absence of 5-FU, the variation frequency of ExoN- rLASV genomic RNA was 2.09-fold of that of wt rLASV, suggesting LASV NP ExoN controls RNA substitution under normal culture conditions. Of note, 5-FU-associated A:G and U:C transitions accounted for 75.7% or 86.1% of the alternate alleles identified in wt or ExoN- rLASV genomic RNA, respectively, following 5-FU treatment (Fig 9). This data confirmed that the nucleotide substitution in 5-FU samples (Fig 7) was largely related to 5-FU incorporation.

For RNA virus, the error-prone RNA replication could increase genetic diversity in virus population and provide benefits for adaptation. However, the mutation rate has to be controlled as even a moderate 1.1-2.8-fold increase in mutation rate could drastically affect virus fitness (50). As negative-sense RNA viruses, how arenaviruses balance the mutation rate while preserving fidelity is largely unknown. Our nucleotide analogue data indicated that LASV NP ExoN contributes to control the rate of viral RNA substitution and ensure the fidelity of viral RNA synthesis. To the best of our knowledge, this is the first example among negative-sense RNA viruses that a non-RdRp viral protein plays a key role in viral RNA fidelity.

In summary, the present study provided evidence that LASV NP ExoN is required for optimal viral RNA replication and mutation control in addition to its role in immune evasion. Loss of LASV NP ExoN activity has multiple impacts on viral RNA replication, including decreased RNA level, increased occurrence of structural deletions in viral RNA, and higher rate of nucleotide substitution. These defects may have more profound impact on viral fitness and pathogenicity *in vivo*. Future studies are warranted to investigate whether NP ExoN of other arenaviruses has similar functions in viral RNA replication. Hemorrhagic fever-causing arenaviruses continue to pose a threat to public health and have pandemic potential. Currently, mutagenic nucleotide analogue T705 (Favipiravir) shows promising activities in animal models of arenavirus infection (51, 52). Our study suggests that targeting NP ExoN may enhance virus sensitivity to nucleotide analogues. Therefore, developing NP ExoN inhibitors could be a valuable strategy to enhance the efficacy of T705 in treatment of LF and other arenavirus-caused hemorrhagic fever diseases.

## Materials and Methods

### Cells and Viruses

Vero cells (CCL-81, ATCC) were maintained in Dulbecco’s modified eagle medium (Hyclone) supplemented with 10% FBS (Invitrogen) and 1% penicillin and streptomycin solution (Hyclone). The LASV (Josiah strain) used in studies was recombinant virus that were rescued using reverse genetic systems in BHK21 cells and passaged only once (P1) in Vero cells as previously described (53). All plasmids used for LASV rescue were confirmed by sequencing. Site-directed mutagenesis on the pmPol I-LASV Sag plasmid was performed using QuikChange II kit (Agilent) according to the manufacturer’s instructions. To construct the rLASV NPD380A mutant, forward primer 5′-CCAAATGCT AAGACCTGGATGGCTATTGAAGGAAGACCTGAAGATC-OH, and reverse primer 5′- GATCTTCAG GTCTTCCTTCAATAGCCATCCAGGTCTTAGCATTTGG-OH were used.

To construct the rLASV NP D389AG392A mutant, forward primer 5′-AAATGCTAAGAC CTGGATGGCTATTGAAGCTAGACCTGAAGATCCAGTGG-OH, and reverse primer 5′- CCACTGGATCTTCAGGTCTAGCTTCAATA GCCATCCAGGTCTTAGCATTT-OH were used. The sequence of mutant rLASV was confirmed by Sanger Sequencing after RT-PCR amplification of viral RNA extracted from infected cells. All infection work with pathogenic arenaviruses was performed at the BSL4 facilities in Galveston National Laboratory in the University of Texas Medical Branch in accordance with institutional health and safety guidelines and federal regulations.

### RNA Extraction, RT-PCR, and real-time RT-qPCR

RNA lysates were prepared using the TRIzol reagent (Life Technology). RNA was purified using the RNeasy Minikit (Qiagen) and treated with DNase I (Qiagen) as previously reported (32, 54). An equal amount of total RNA (0.5 to 1µg) was reverse transcribed to cDNA at 55 °C for 60 minutes using high-fidelity reverse transcriptase SuperScript IV (Invitrogen) per manufacturer’s instruction. The high-fidelity reverse transcriptase SuperScript IV has low RNase H activity and is suitable for long cDNA synthesis. Random hexamer primers and sequence specific primers were used in RT as specified in each experiment. cDNA samples were treated with RNase H. In RT of S gRNA, an S gRNA 3′UTR specific primer (5′-CGCACAGTGGATCCTAGGCTA-OH) was used. In RT of L gRNA, an L gRNA 3′UTR specific primer (5′- CACCGAGGATCCTAGGCATTAAGGCTATC-OH) was used in cDNA synthesis. PCR amplification of close to full-length Sg cDNA (3391bp) was conducted with Platinum SuperFi DNA Polymerase (Invitrogen) using forward primer 5′-ATCCTAGGCATTTTTGGTTGC-OH and reverse primer 5′-CGCACAGTGGATCCTAGGCTA-OH. PCR amplification of close to full-length Lg cDNA (7265 bp) was performed with Platinum SuperFi DNA Polymerase using forward primer 5′-CACCGAGGATCCTAGG CATTAAGGCTATC-OH and reverse primer 5′-ATCCTAGGCAATTTGGTTGTTCTTTTTTGAG-OH. Real-time quantitative PCR (qPCR) was performed with SsoAdvanced Universal SYBR green Supermix (BioRad) on a CFX96 real-time PCR detection system (BioRad) as described previously (32, 54). To measure the viral RNA level at NP locus and L locus, cDNA synthesized with random primers was used in a qPCR assay. LASV NP forward primer (5′-GAAGGGCCT GGGAAAACACT-OH) and LASV NP reverse primer (5′-AGGTAAGCCCAGCGGTAAA

C-OH) were used for the NP locus. LASV L forward primer (5′-CAGCAGGTCAGACGA AGTGT-OH) and LASV L reverse primer (5′-GTTGTGCATAGGGGAGGCTT-OH) were used for the L locus. The cDNA synthesized with SgRNA- and LgRNA-specific primers as indicated above was used in a qPCR assay to measure the level of LASV Sg RNA and Lg RNA. The qPCR data was analyzed with the CFX Manager software (BioRad). The RNA level of each target gene was normalized to that of the housekeeping gene β- actin with validated primers from BioRad. All experiments were performed separately in triplicates.

### PacBio SMRT long-read sequencing of LASV SgRNA

Vero cells was infected by ExoN- rLASV and wt rLASV at a MOI of 1. At 48 hpi, total RNA was purified from infected cells. RT-PCR amplification of the close-to-full-length SgRNA was performed as described above. PCR amplicons (approximately 3.4 kb) were gel purified using Monarch DNA gel extraction kit (NEB). Library construction, PacBio SMRT long-read sequencing and CCS determination was performed by Azenta Life Sciences (USA). At least 1 µg of amplicon was ligated to barcoded adapters (Pacific Biosciences) for SMRTbell library construction and then sequenced on the PacBio Sequel IIe platform with v3.0 chemistry. Raw reads with a minimum number of passes greater than 3 were used to generate the CCS reads using PacBio SMRTLINK v.10.1. HiFi reads (>=99% accuracy) were extracted. Data analysis was performed utilizing *Galaxy*: an open source, web-based platform supported by NIH, NSF, and the Texas Advanced Computing Center. CCS reads were filtered for those with size 3100- 3394bp and high-quality reads (Phred>20) using Filter FASTQ and mapped to the LASV reference seq (MH358389) with *Minimap2* (PacBio HiFi reads vs reference mapping (- k19 -w19 -U50,500 -g10k -A1 -B4 -O6,26 -E2,1 -s200)). Variants with cutoff >3% frequency were called with *iVar* and visualized using the Integrated Genomics Viewer (IGV). The raw sequencing data has been deposited in Sequence Read Archive (SRA) databases hosted by the National Library of Medicine’s National Center for Biotechnology Information (NCBI), NIH.

### Mutagenic nucleotide analogue treatment

5-fluorouracil (Cat# 03738, Sigma) and N4-hydroxycytidine (EIDD-1931) (Cat#9002958, Cayman) were dissolved in DMSO (cell culture grade, Sigma) and further diluted in DMEM media contain 2% FBS and 0.1% DMSO. Vero cells were mock-treated or treated with 5-FU or EIDD-1931 at different concentrations as indicated in each experiment for 4 hr and then infected with wt rLASV and ExoN- rLASV at a MOI of 0.1 for 1 hr. Virus inoculums were removed and replaced with fresh media (DMEM + 2% FBS and 0.1% DMSO) containing 5-FU or EIDD-1931 of different concentrations as indicated. At 48 hpi, the supernatants of infected cells were harvested and subjected to plaque assay to determine virus titers. The viability of Vero cells treated with 5-FU and EIDD-1931 alone were assessed using CellTiter-Glo Viability Assay (Promega).

### Illumina NGS analysis of LASV RNA amplicons following 5-FU treatment

Virus infection and 5-FU treatment of Vero cells were performed as described above. The supernatants of virus infected cultures were harvested and clarified by centrifugation at 3,000 rpm to remove cell debris. Virions were concentrated and purified with Amicon Ultra-15 Centrifugal Filter Units (Millipore, MW cutoff 100 kD). RNA extraction and reverse transcription with S gRNA and L gRNA-specific primers were performed as described above. PCR were performed with high-fidelity Platinum SuperFi DNA Polymerase. For S gRNA, two overlapping amplicons were amplified. For L gRNA, three overlapping amplicons were amplified. The primers used in amplicon preparation are listed in supplementary information (S2 Table). All amplicons were purified from agarose gel after electrophoresis. Library construction, Illumina deep sequencing (Hiseq, 2×150 bp, pair-end) and raw data process was performed by Azenta Life Sciences (USA). The obtained reads had a mean read quality score over 37. More than 90% of the bases had a Phred quality score greater that 30 (i.e., sequencing accuracy of 99.9%). Data processing and analysis was performed on *Galaxy*, a web-based platform supported by NIH, NSF, and the Texas Advanced Computing Center. The raw data (in FASTQ format) was trimmed with *Trimmomatic* (55) to remove adaptor sequences, and then mapped to the LASV reference seq (MH358389) with minimap2 (41) (short reads without splicing (-k21 -w11 --sr -F800 -A2 -B8 -O12,32 -E2,1 -r50 -p.5 -N20 -f1000,5000 -n2 -m20 -s40 -g200 -2K50m --heap-sort=yes --secondary=no). Read depth was assessed with Samtools (56). The read depth was higher than 1,000,000 reads at each site on LASV genomic RNA, except for some positions at IGR. Low frequency variants were called with LoFreq (57) (minimal base calling quality>20 for reference bases and alternate bases, minimum mapping quality 20) from one million reads at each position. Single nucleotide variations with higher than 0.1% frequency was included for analysis to minimize background noise due to sequencing error. To ensure the confidence of data quality, reads with strand bias greater than 100 (SB>100) were excluded. The data were imported to Excel for mutation analysis. Raw NGS data have been deposited in SRA databases hosted by NCBI, NIH.

## Acknowledgments

This work was supported in part by Public Health Service grants 1R21AI166985 (to CH) R01AI093445 (to SP) and R01AI129198 (to SP). CH was also supported by Institute for Human Infections & Immunity, UTMB Pilot Study Fund (P86035) and would like to acknowledge Galveston National Laboratory (supported by the Public Health Service award 5UC7AI094660) for support of research activity. Research work in Dr. Paessler’s lab was also supported by the John. S. Dunn Distinguished Chair in Biodefense endowment. E.K.M was supported by National Institutes of Health T32 training grant AI060549.

---

S1 Fig: Read depth of viral genomic RNA in NGS analysis.

S2 Table: Primers used in PCR amplification of LASV amplicon for NGS analysis.

