## Supplementary figures and images for "Lassa virus NP DEDDh 3′-5′ exoribonuclease activity is required for optimal viral RNA replication and mutation control"

### Supplemental Figure

#### S1 Fig

A. Read depth of S RNA

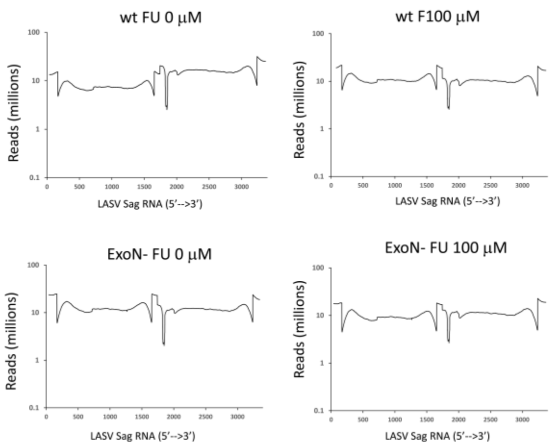


B. Read depth of L RNA

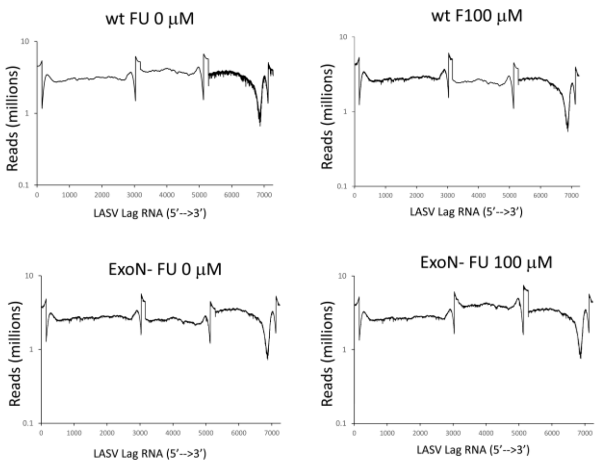
