## Supplemental Table for "Lassa virus NP DEDDh 3′-5′ exoribonuclease activity is required for optimal viral RNA replication and mutation control"

S2 Table: Primers used in PCR amplification of LASV amplicon for NGS analysis

| Primer | Sequence |
|---|---|
| S1 amplicon Forward | 5'-AACGACTCTAGGTGTCGATGTTCT |
| S1 amplicon Reverse | 5'-CTATTGGATTGCGCTTTGCT |
| S2 amplicon Forward | 5'-CCTAGGCATTTTTGGTTGCGC |
| S2 amplicon Reverse | 5'-TCAGGGGTCCGATGACATAAGG |
| L1 amplicon Forward | 5'-ATCCTAGGCAATTTGGTTGTCTTTTTTTGAG |
| L1 amplicon Reverse | 5'-CATCTGAGTCTGACCTTGAGTATTCTTGG |
| L2 amplicon Forward | 5'-CCTAAGACCCATGCACCCAGT |
| L2 amplicon Reverse | 5'-CTTCTCAAGGAAAGGAAAGTACCTTCTCA |
| L3 amplicon Forward | 5'-GGACACTGTGACATATGTCCACAGT |
| L3 amplicon Reverse | 5'-CACCGAGGATCCTAGGCATTAAGG |
